# Testing the behavioral origins of novelty: did increased aggression lead to scale-eating in pupfishes?

**DOI:** 10.1101/280123

**Authors:** Michelle E. St. John, Joseph A. McGirr, Christopher H. Martin

## Abstract

How novelty evolves is still largely unknown. Environmental changes are often assumed to precede novelty; however, behavioral shifts may also play a role. Here, we examine whether a shift in aggression explains the origin of a novel scale-eating pupfish species (*Cyprinodon desquamator*) within an adaptive radiation on San Salvador Island, Bahamas. We compared aggression using behavioral and gene expression data across three sympatric species in the San Salvador radiation (generalist, snail-eating specialist, and scale-eating specialist), and additionally measured behavioral aggression in an outgroup generalist from North Carolina. Surprisingly, we found increased behavioral aggression and differential expression of aggression-related genes in both the scale-eating and snail-eating species. Furthermore, male scale-eaters and female snail-eaters showed the highest levels of aggression compared to other groups. Differential gene expression in each specialist during larval development also suggested sex-mediated differences in male-male aggression and maternal care. Ultimately, our data indicate that aggression is not unique to scale-eating specialists. Instead, selection may increase aggression in other contexts such as niche specialization, mate competition, or selection on other ecologically relevant traits, including jaw size. Indeed, some adaptive variants associated with oral jaw size in the San Salvador radiation occur in genetic pathways with pleiotropic effects on aggression.

## Introduction

The origins of evolutionary novelty are still poorly understood. For example, both novel behavior and morphology play a role in the evolution of novel resource use, but their relative importance and order in which they evolve are still unknown. Changes in behavior may precede the evolution of novel morphologies, as they can expose organisms to novel environments and selective pressures (Huey et al. 2003). Investigations of novelty, however, overwhelmingly ignore this possibility (although see: Huey et al. 2003; Losos et al. 2004; Duckworth 2006). Instead, previous studies have focused on novel adaptive morphologies or on how environmental changes expose organisms to new selective pressures (Liem 1973; Barton and Partridge 2000; Janovetz 2005a; Hulsey et al. 2008). One reason behavior has been overlooked as an origin for novelty is because it is still unclear whether it drives or inhibits evolution, as behavior is often extremely plastic. However, in order to determine if novelty has a behavioral origin we must first understand its variation within and among taxa.

One outstanding example of novelty is lepidophagy (scale-eating) in fishes. Scale-eating has been documented in five freshwater and seven saltwater families of fishes and has independently evolved at least 19 times (Sazima 1983; Janovetz 2005b; Martin and Feinstein 2014; Kolmann et al. 2018). Scale-eating includes both novel morphologies and novel behaviors. For example, some scale-eaters have premaxillary external teeth for scraping scales (Novakowski et al. 2004), some use aggressive mimicry to secure their prey (Boileau et al. 2015), others sneak scales from the surface of fish that they are cleaning (Losey 1979), and still others use ambush tactics to obtain scales (Nshombo et al. 1985). Even though scale-eating is an outstanding example of the convergent evolution of novel trophic ecology across disparate environments and taxa and displays a wide variety of morphologies and behaviors, its origins have yet to be explored.

There are currently only three hypothesized behavioral origins for scale-eating. The algae-grazer hypothesis predicts that scale-eating arises from the incidental ingestion of scales during algae scraping (Fryer et al. 1955; Greenwood 1965; Sazima 1983). Many scale-eaters are closely related to algae-grazers. For example, many rock-dwelling Malawi cichlids are algae-scrapers (Greenwood 1965; Fryer and Iles 1972; Ribbink et al. 1983), but the radiation also includes two sister species of scale-eaters (*Corematodus shiranus* and *Corematodus taeniatus*) and a second independent origin of scale-eating in *Genyochromis mento* (Trewavas 1947; Greenwood 1965). Similarly, the extinct Lake Victorian scale-eater *Haplochromis welcommei* was nested within rock-dwelling algae scrapers (Greenwood 1965). This hypothesis, however, does not address why algae-grazing fish would seek food on the surface of other fish (Greenwood 1965). The second hypothesis, termed the cleaner hypothesis, tries to address this gap by arguing that scale-eating arose from the incidental ingestion of scales during the consumption of ectoparasites from the surface of other fishes (Greenwood 1965; Sazima 1983). One line of evidence supporting this hypothesis is that cleaner fish sometimes eat scales. For example, the Hawaiian cleaner wrasse (*Labroides phthirophagus*) and two species of juvenile sea chub (*Hermosilla azurea* and *Girella nigricans*) consume both ectoparasites and scales (Demartini and Coyer 1981; Sazima 1983; Losey 1972). However, most scale-eating fishes are not known to forage on ectoparasites, nor are they closely related to fish that do. In fact, the closest examples of this are the false cleaner fishes (*Aspidontus taeniatus* and *Plagiotremus rhinorhynchs*) who aggressively mimic cleaner wrasse (*Labroides dimidiatus*) in order to consume scales. Despite their morphological similarities, however, these fish are not closely related (Hundt et al. 2014). Finally, the aggression hypothesis predicts that scale-eating evolved due to the accidental ingestion of scales during inter- or intra-species aggression (Sazima 1983). This hypothesis is supported by the fact that two characid species of scale-eaters (*Probolodus heterostomus* and *Exodon paradoxus*) are closely related to the aggressive *Astyanax* tetras (Sazima 1983; Kolmann et al. 2018); a similar argument can be made for the scale-eating piranha (*Cataprion mento*) (Janovetz 2005).

The scale-eating pupfish, *Cyprinodon desquamator*, is an excellent species for investigating the origins of scale-eating because it is, by far, the youngest scale-eating specialist. The species is nested within a sympatric adaptive radiation of pupfishes endemic to the hypersaline lakes of San Salvador island, Bahamas (Martin and Wainwright 2011, 2013a). In addition to the scale-eating pupfish, this radiation also includes a widespread generalist (*C. variegatus)* and an endemic snail-eating specialist (*C. brontotheroides).* Other generalist pupfish lineages are also distributed across the Caribbean and western Atlantic Ocean. Phylogenetic evidence indicates that scale-eater’s most recent ancestor was a generalist feeder (Martin 2016; Richards and Martin 2017). Furthermore, geological evidence suggests that the hypersaline lakes of San Salvador island—and thus the radiation containing the scale-eater—is less than 10 thousand years old (Hagey and Mylroie 1995; Martin and Wainwright 2013a,b).

We investigated the possible behavioral origins of novelty by examining whether a shift in aggression led to the evolution of scale-eating in pupfish. We compared measures of aggression using both behavioral and gene expression data between scale-eaters and closely related species within their radiation (a sympatric generalist species and a snail-eating species from San Salvador Island) as well as an additional outgroup (a generalist species from North Carolina). We predicted high levels of aggression in scale-eaters, intermediate levels in the San Salvador generalist and snail-eating pupfish, and low levels of aggression in the North Carolina (NC) outgroup generalist. Surprisingly, we found that male scale-eaters and female snail-eaters displayed increased levels of aggression, while other groups displayed relatively low levels of aggression. This suggests that aggression alone cannot explain the origins of scale-eating in pupfish. We also identified promising candidate genes that are differentially expressed between species associated with differences in aggression and linking behavior, morphology, and the evolution of novel ecology.

## Methods

### Sampling

Generalist, snail-eating, and scale-eating pupfish were collected by seine from Crescent Pond, Great Lake, Little Lake, Osprey Lake, and Oyster Pond of San Salvador Island, Bahamas in July, 2016. In June 2017, generalist pupfish were collected by seine from the Cape Fear river drainage (Fort Fisher) on the coast of North Carolina. Fishes were housed in 40 – 80 liter tanks in mixed-sex groups at 5-10 ppt salinity in temperatures ranging from 23°C-30°C. Fish were fed a diet of frozen blood worms, frozen mysis shrimp, or commercial pellet food daily.

### Aggression assay

We quantified levels of aggression for each pupfish species and sex using mirror tests (Vøllestad and Quinn 2003; Francis 2010). To control for individual size and motivation, we incited aggression using a compact mirror (10 cm X 14 cm) placed in a 2-liter trial tank (25 cm X 16 cm X 17 cm). We randomly chose adult fish and isolated each one in 2-liter tanks that contained a single bottom synthetic yarn mop for cover and opaque barriers between adjacent tanks. We gave the fish at least 12 hours to acclimate to their new environment before performing an assay. During a 5-minute focal observation period, we measured three metrics as a proxy for aggression: latency to approach mirror image, latency to attack mirror image, and total number of attacks toward the mirror image. A trial began as soon as the mirror was securely lowered into the tank. We measured latency to approach as the time elapsed before an individual approached the mirror to within one-body length. Similarly, we measured latency to attack as the time elapsed before an individual attacked their mirror image for the first time. Finally, we counted the total number of attacks an individual performed during the entirety of the trial. We also measured the standard length of each fish after the trial.

### Statistical analyses

We used time-to-event analyses to determine if species and sex were associated with 1) latency to approach their mirror image and 2) latency to attack their mirror image. For the latency to approach metric (time in seconds) and the latency to attack metric (time in seconds) we used a mixed-effects Cox proportional hazards model (coxme package; Therneau 2012) in R (R Development Core Team 2016). These models allow right censored data, i.e. individuals who did not approach or attack their mirror image were not excluded and contributed to Kaplan-Meier estimates and time-to-event curves (Rich et al. 2010). For both the latency to approach model and the latency to attack model we included species, sex, and their interaction as fixed effects and lake population as a random effect. We compared these models to equivalent models that also included size (log scale) as a covariate using AICc (Burnham and Anderson 2002; stats package; R Development Core Team 2016). For the latency to approach model, size was non-significant (*P* = 0.36) and we removed it from the model. For the latency to attack model, however, size was a significant covariate and retained in the final model. We used the likelihood ratio test to determine if species, sex, or their interaction were associated with latency to approach or attack the mirror image. Additionally, we used a Cox proportional hazards model without mixed effects to plot the resulting time-to-event curves and made pairwise comparisons between curves using log-rank tests (Survival Package; survminer package; Therneau 2015).

We analyzed the total number of attacks using a generalized linear mixed model (GLMM) with a negative binomial distribution for this response variable. We modeled species, sex, and their interaction as fixed effects, and population as a random effect. We compared this model to a model including size (log scale) as a continuous covariate using AICc, but accounting for size did not substantially increase the likelihood of the model (ΔAICc = 0.96). We used a Wald chi-square test (type II) to determine if species, sex, or their interaction significantly affected the total number of attacks performed and used Tukey’s HSD to make direct comparisons between groups.

### Identifying candidate genes affecting differences in aggression between species

We searched a previously published dataset of 15 San Salvador pupfish transcriptomes to identify candidate genes underlying behavioral differences in aggression among all three species (Mcgirr and Martin 2018). This previous study did not analyze gene expression pathways annotated for effects on behavior. Briefly, purebred F_1_ and F_2_ offspring from the three-species found on San Salvador island were raised in a common garden laboratory environment. Larvae were euthanized in an overdose of MS-222 at 8-10 days post fertilization (dpf) and were immediately preserved in RNAlater (Ambion, Inc.) and stored at −20 C after 24 hours at 4 C. Total mRNA was extracted from 6 generalists, 6 snail-eaters, and 3 scale-eaters (RNeasy kits, Qiagen). Stranded sequencing on an Illumina HiSeq 4000 at the High Throughput Genomic Sequencing Facility at UNC Chapel Hill resulted in 363 million raw reads that were aligned to the *Cyprinodon variegatus* reference genome (NCBI, C. *variegatus* Annotation Release 100, total sequence length =1,035,184,475; number of scaffold=9,259, scaffold N50, =835,301; contig N50=20,803; Lencer et al. 2017). Aligned reads were mapped to annotated features using STAR (v. 2.5(33)), with an average read depth of 309x per individual and read counts were generated across annotated features using the featureCounts function from the Rsubread package (Liao et al. 2013). DEseq2 (Love et al. 2014, v. 3.5) was used to normalize counts and identify: 1) 1,014 differentially expressed genes between snail-eaters *vs* generalists and 2) 5,982 differentially expressed genes between scale-eaters *vs* generalists (McGirr and Martin 2018).

We identified one-way best hit zebrafish orthologs for genes differentially expressed between 1) snail-eaters *vs* generalists (n=722) and 2) scale-eaters *vs* generalists (n=3,966) using BlastP (V. 2.6; E-value <1). We compared this list of orthologs to gene ontologies describing aggressive behavior (GO: 0002118), inter-male aggressive behavior (GO: 0002121), maternal aggressive behavior (GO:0002125), maternal care behavior (GO: 0042711), and territorial aggressive behavior (GO: 0002124; AmiGo; Carbon et al. 2009; Ashburner et al. 2000; The Gene Ontology Consortium 2017). We also searched gene ontologies for three hormone pathways commonly associated with aggression (the vasopressin pathway, the androgen pathway, and the estradiol pathway).

## Results

### Behavioral aggression

Male scale-eaters and female snail-eaters showed increased levels of aggression compared to other groups. Scale-eaters (at the species level) approached their mirror image significantly more than NC generalists (Table 1A; Fig. 1A; log-rank test, *P* = .038). Additionally, male scale-eaters attacked their mirror image significantly more than male San Salvador generalists (Table 1B; Fig. 1B; log-rank test, *P* =0.032). Lastly, male scale-eaters were the only group to exhibit increased aggression compared to their female counterparts. More male scale-eaters attacked their mirror image than did female scale-eaters (Table 1B; Fig. 1B: log-rank test, *P* = 0.003), and they performed significantly more total attacks (Table 1C; Fig. 2; Tukey HSD, *P* = 0.0003).

**Table 1.**
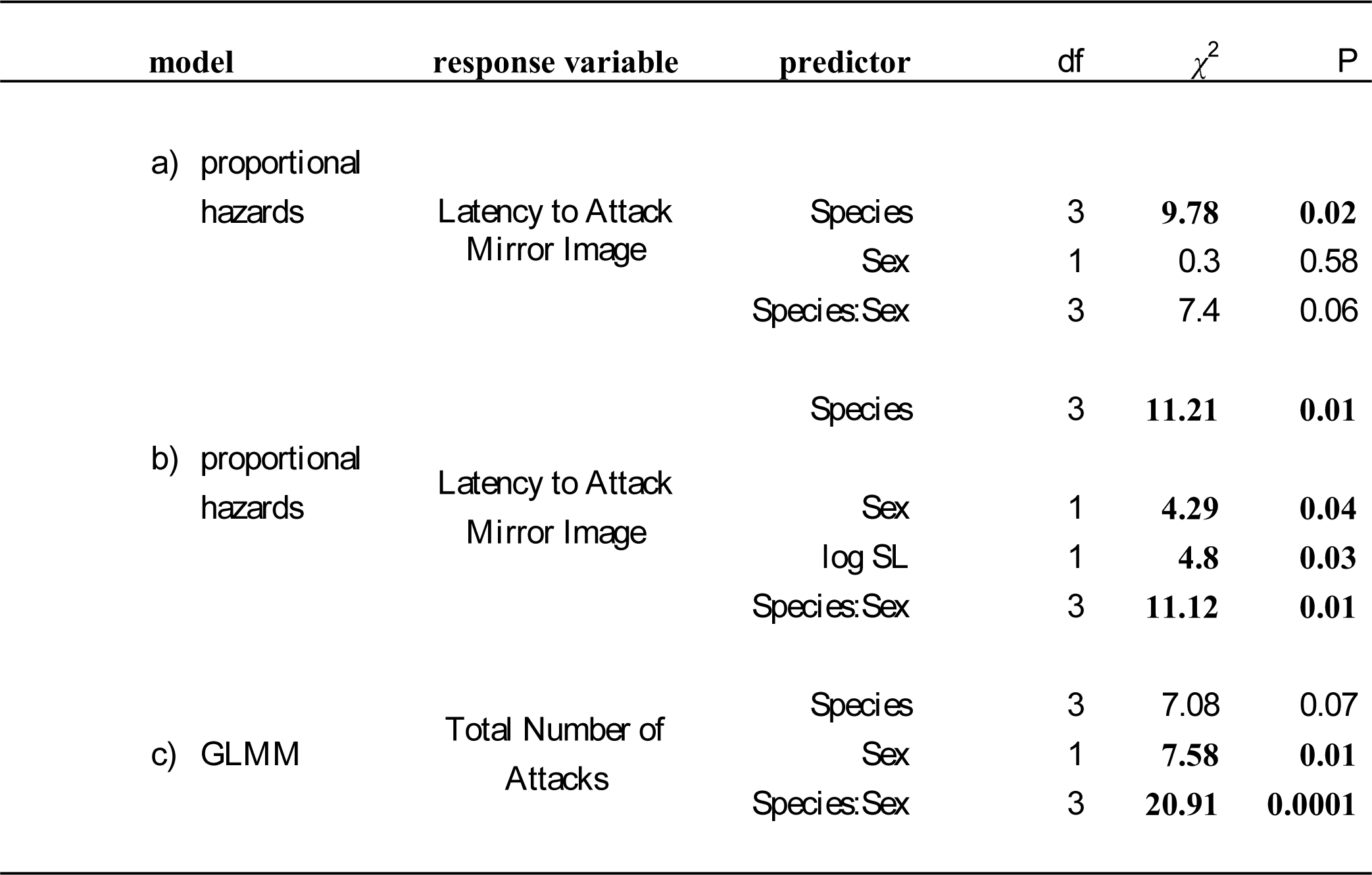
Results of likelihood ratio test for: a) the latency to approach mirror image (mixed-effect Cox proportional hazards model); b) the latency to attack mirror image (mixed-effect Cox proportional hazards model); and c) the total number of attacks (generalized linear mixed model). Significant results are indicated in bold.

**Figure 1.**
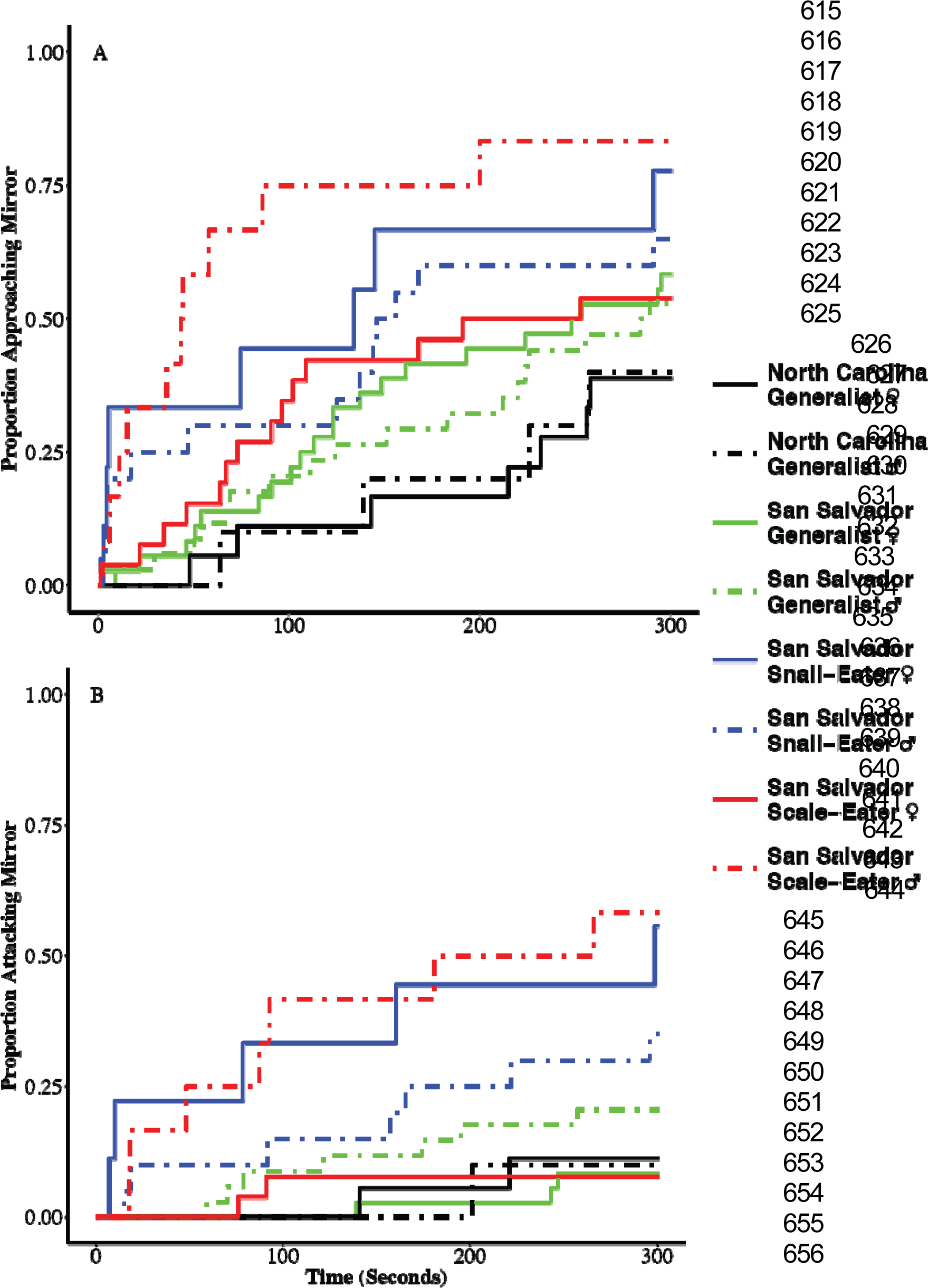
Time-to-event curves for *a)* the latency to approach mirror image (Cox proportional hazards model) and *b)* the latency to attack mirror image (Cox proportional hazards model).

**Figure 2.**
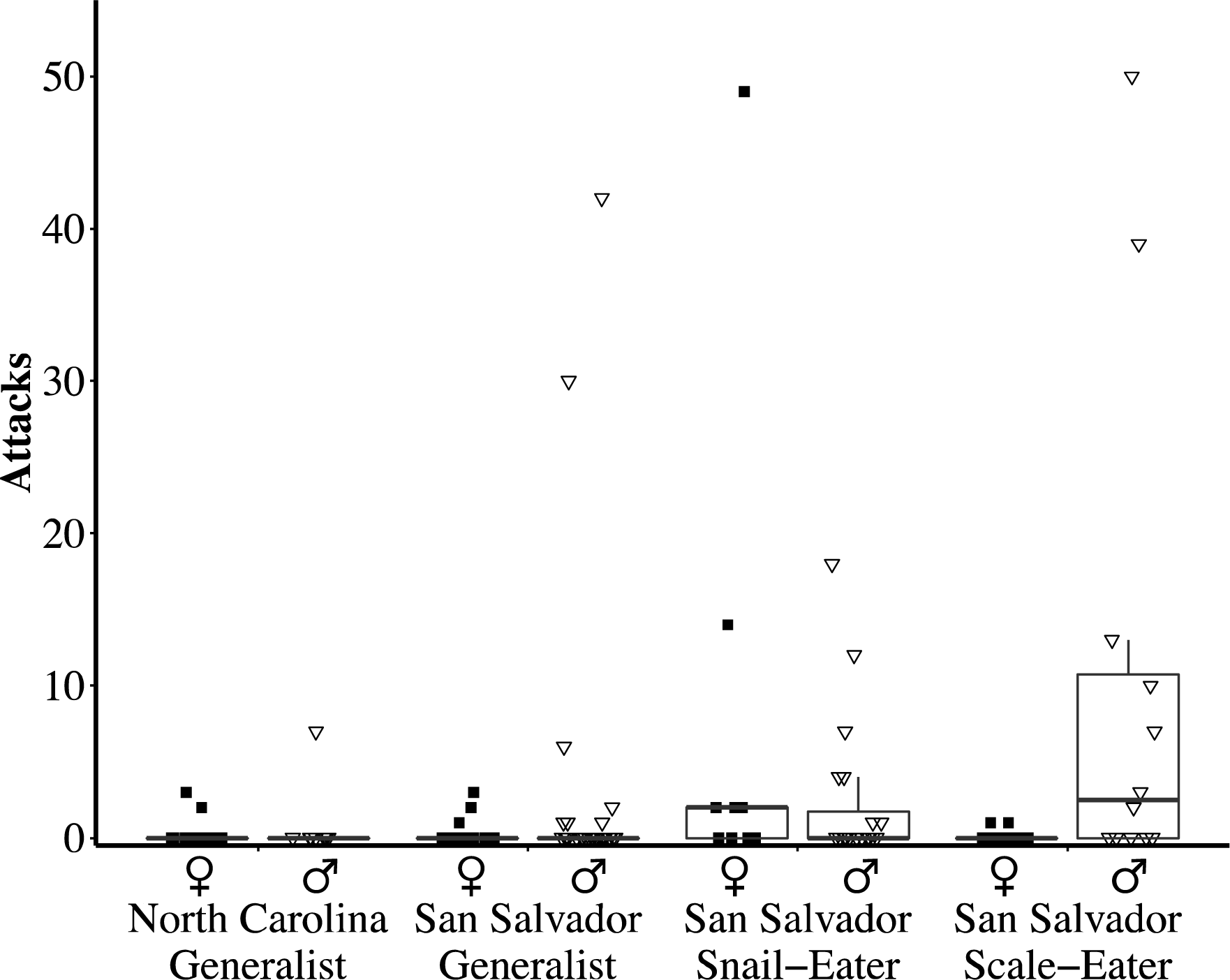
Box plots illustrating the total number of attacks performed by each species and sex (*n* = 165 total individuals tested). Squares represent the total number of attacks for individual females, while triangles represent the total number of attacks for individual males.

Female snail-eaters showed a similar pattern of increased aggression. Snail-eaters (at the species level) approached their mirror image significantly more than NC generalists (Table 1A; Fig. 1A; log-rank test, *P* = .038). Significantly more female snail-eaters attacked their mirror image than all other female groups (Table 1B; Fig. 1B; log-rank test, NC generalist *P* = 0.032, San Salvador generalist *P* = 0.0027, San Salvador scale-eater *P* = 0.0081). Finally, female snail-eaters performed significantly more attacks than female San Salvador scale-eaters and generalists (Table 1C; Fig. 2; Tukey HSD, generalist *P* = 0.0032, scale-eater *P* = 0.0006).

### Gene Expression

We searched genes that were differentially expressed between scale-eaters *vs* generalists or between snail-eaters *vs* generalists for gene ontologies describing aggressive behavior, inter-male aggressive behavior, maternal aggressive behavior, maternal care behavior, the vasopressin hormone pathway, and the androgen hormone pathway (Table 2). Despite over one thousand differentially expressed genes at this developmental stage, only five genes were associated with these aggression-related ontologies in the snail-eater *vs* generalist comparison (Table 2A). Scale-eaters also exhibited differential expression of genes associated with inter-male aggression and vasopressin when compared to their generalist sister species (Table 2B). Interestingly, one of these genes (*gnaq*) contains a fixed variant in scale-eaters, which is known to function in craniofacial development and shows signs of a hard selective sweep (McGirr and Martin 2017; McGirr JA unpublished data). None of these ontologies were significantly over-represented in either species comparison, which were instead enriched for cranial skeleton, metabolism, and pigmentation terms (McGirr and Martin 2018).

**Table 2.**
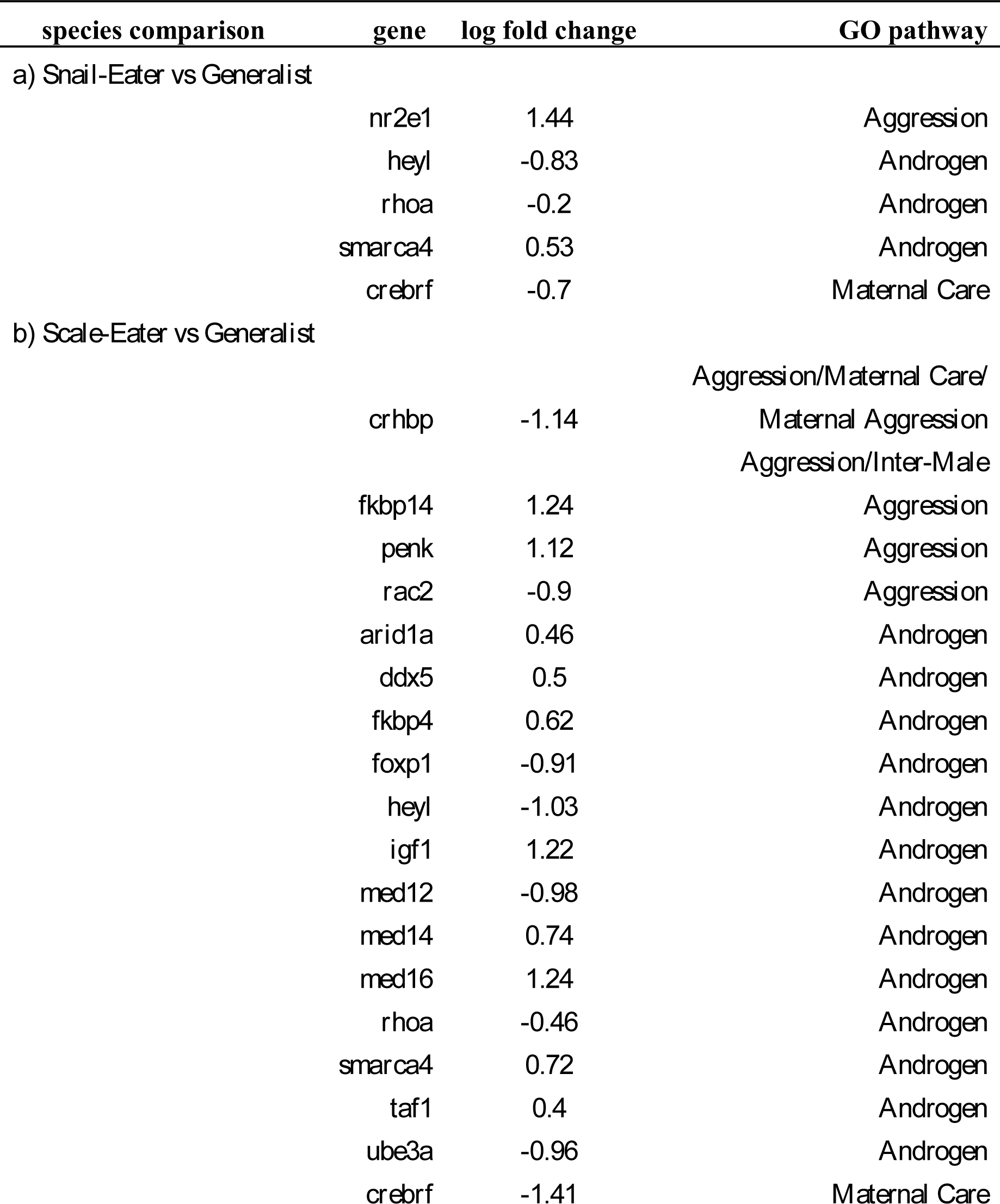
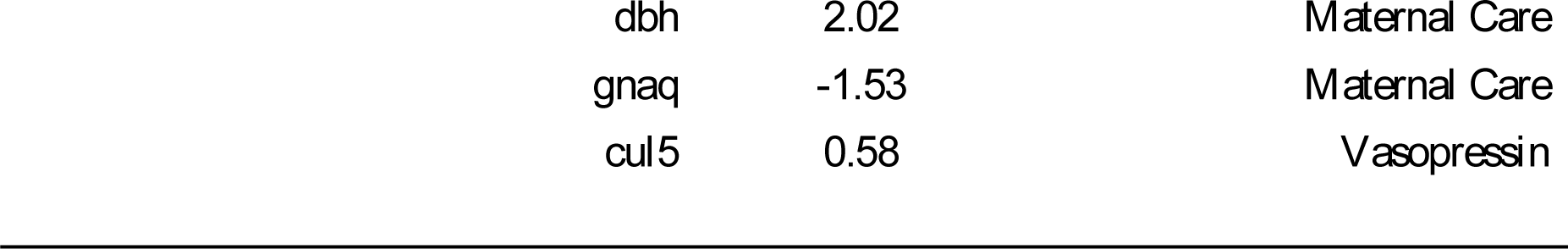
List of all differentially expressed genes in aggression-related pathways between *a)* sail-eaters *vs* generalists and *b)* scale-eaters *vs* generalists. 3-6 individuals of each species sampled at 8-10 dpf.

| species comparison | gene | log fold change | GO pathway |
| --- | --- | --- | --- |
| *a) Snail-Eater vs Generalist* |  |  |  |
|  | *nr2e1* | 1.44 | Aggression |
|  | *heyl* | -0.83 | Androgen |
|  | *rhoa* | -0.2 | Androgen |
|  | *smarca4* | 0.53 | Androgen |
|  | *crebrf* | -0.7 | Maternal Care |
| *b) Scale-Eater vs Generalist* |  |  |  |
|  | *crhbp* | -1.14 | Aggression/Maternal Care/<br>Maternal Aggression<br>Aggression/Inter-Male |
|  | *fkbp14* | 1.24 | Aggression |
|  | *penk* | 1.12 | Aggression |
|  | *rac2* | -0.9 | Aggression |
|  | *arid1a* | 0.46 | Androgen |
|  | *ddx5* | 0.5 | Androgen |
|  | *fkbp4* | 0.62 | Androgen |
|  | *foxp1* | -0.91 | Androgen |
|  | *heyl* | -1.03 | Androgen |
|  | *igf1* | 1.22 | Androgen |
|  | *med12* | -0.98 | Androgen |
|  | *med14* | 0.74 | Androgen |
|  | *med16* | 1.24 | Androgen |
|  | *rhoa* | -0.46 | Androgen |
|  | *smarca4* | 0.72 | Androgen |
|  | *taf1* | 0.4 | Androgen |
|  | *ube3a* | -0.96 | Androgen |
|  | *crebrf* | -1.41 | Maternal Care |

|  |  |  |
| --- | --- | --- |
| *dbh* | 2.02 | Maternal Care |
| *gnaq* | -1.53 | Maternal Care |
| *cul5* | 0.58 | Vasopressin |

## Discussion

The origins of novelty have overwhelmingly been examined from a morphological perspective, often ignoring behavior’s potential role. However, shifts in behavior may also be a viable origin for novelty. Unfortunately, only a few previous studies have directly investigated this possibility. The origin of novelty in the Pacific field cricket (*Teleogryllus oceanicus*)—which exhibits a novel silent morph—is one of the few examples of evolutionary novelty with a behavioral origin (Zuk et al. 2006; Tinghitella and Zuk 2009; Bailey et al. 2010). Increased brain size in birds has also been linked to behavioral shifts and novelty. Birds that display innovative feeding behaviors have larger brains and are more successful at invading novel environments (Nicolakakis and Lefebvre 2000; Sol and Lefebvre 2000; Overington et al. 2009). Likewise, the role of behavior in evolutionary novelty has also been explored in western bluebirds *(Sialia Mexicana;* Duckworth 2006) and *Anolis* lizards (Losos et al. 2004, 2006). Despite the growing empirical evidence of behavior’s role in evolutionary innovation, a consensus has not yet been reached on whether behavior ultimately drives or inhibits novelty. Furthermore, studies that investigate behavioral origins of novelty rarely do so using both behavioral and genetic approaches. However, by leveraging our gene expression data, we gained some mechanistic insight into the divergent origins of increased behavioral aggression in each specialist species.

We tested whether increased aggression contributed to the origin of scale-eating in a species of Caribbean pupfish using both behavioral and gene expression data. The aggression hypothesis predicts that scale-eating arose due to increased inter- and intra-specific aggression (Sazima 1983). Contrary to these predictions, both snail-eaters and scale-eaters showed increased levels of aggression. Our gene expression data supported these findings, as both scale-eaters and snail-eaters showed differential expression of genes involved in several aggression-related pathways during larval development. Additionally, both scale-eaters and snail-eaters displayed surprising differences in aggression between the sexes. While male scale-eaters showed increased levels of aggression, female scale-eaters showed extremely low levels of aggression. Conversely, female snail-eaters showed increased levels of aggression compared to females of other species. These results suggest that the aggression hypothesis alone cannot explain the evolution of scale-eating. Instead, selection may have favored increased levels of aggression in other contexts, such as mate competition or trophic specialization in general. Increased levels of aggression could have also arisen indirectly due to selection for other behaviors or traits, including several genes involved in both aggression and craniofacial morphology (e.g. *gnaq*).

One caveat is that there is still discussion whether mirror tests accurately predict levels of aggression in the field. Balzarini et al. (2014) argue that, while mirror tests are a valid method of measuring aggression in some species, they are inappropriate for others. For example, some species use lateral displays of aggression which primarily occur head to tail—a maneuver that is impossible with a mirror image. Additional studies also indicate that mirror tests may not accurately predict aggressive display frequency, duration, or orientation (Elwood et al. 2014; Arnott et al. 2016). It is possible that our method of measuring aggression may have underestimated aggression for scale-eating pupfish. This may be particularly true for female scale-eaters. Our study primarily measured direct displays of aggression (i.e. attacks), however, females often display aggression indirectly (Rosvall 2011; Stockley and Campbell 2013). Our methods of measuring aggression, therefore, may have missed increased levels of aggression in female scale-eaters while still detecting them in males.

A second caveat is that we compared differential gene expression in an early larval developmental stage, 8-10 dpf, long before sexual maturity in this species. Thus, we are not comparing adult differences in gene expression between the sexes in each species. Instead, by examining early larval stages our gene expression analyses provide insight into species-specific differences in aggression-related genetic pathways established during an early developmental timepoint. This has the advantage of defining structural developmental differences in each species, rather than transient differences in transcription between adult male and female fish sensitive to dominance status, reproductive state, and mood. Furthermore, we found surprising congruence between our behavioral and transcriptomic data supporting the conclusions of increased aggression in both San Salvador specialists due to different aggression-related genetic pathways.

### New hypotheses for varying levels of aggression between pupfish species

If increased levels of aggression are not associated with scale-eating, then what explains this variation between species? One possibility is that selection may have directly favored increased aggression in the context of dietary specialization. Aggression may be positively correlated with traits associated with specialization (Genner et al. 1999; Peiman and Robinson 2010; Blowes et al. 2013), suggesting that specialists should show increased levels of aggression compared to generalists. Existing evidence supports this as increased levels of aggression have been documented in specialist butterfly fish (*Chaetodontids;* Blowes et al. 2013), specialist striped surfperch (*Embiotoca lateralis*; Holbrook and Schmitt 1992), and have even been observed in game-theoretic simulation models (Chubaty et al. 2014). A second possibility is that increased aggression may be associated with colonizing a novel niche. Aggression is tightly correlated with boldness in a phenomenon termed the aggressiveness-boldness syndrome (Sih et al. 2004). Many studies have shown that increased boldness in species such as cane toads, mosquitofish, and Trinidadian killifish leads to increased dispersal into novel habitats or niches (Fraser et al. 2001; Rehage and Sih 2004; Gruber et al. 2017). This relationship indicates that increased aggression may be an incidental effect of selection for increased boldness and occupation of a novel niche. Our data supports either scenario, as we observed increased aggression in both San Salvador specialists. Neither of these hypotheses, however, explain the variation in aggression between sexes.

### Aggression and mating system

Increased aggression may be favored in the context of courtship or mate competition. It is well documented across multiple taxa that the sex with the higher potential reproductive rate should have increased levels of aggression as they must compete more heavily for access to mates (Clutton-Brock and Parker 1992; Andersson 1994; Jennions and Petrie 2007). Normally, males have higher potential reproductive rates since mating is energetically cheap for them (Trivers 1972). Scale-eaters, and *Cyprinodon* pupfishes in general, seem to adhere to this standard as they mate in a lekking system and do not provide parental care (Gumm 2012). Male scale-eaters may be more aggressive to compete for mates. We found some support for this in our gene expression data. Specifically, we found differential expression in the *rac2* and *ube3a* genes between scale-eaters *vs* generalists. The *rac2* gene is associated with the visualization of visible light, metabolism, and behavior (Elsaesser et al. 2010; Goergen et al. 2014). Mutations in the *rac2* gene affect both male aggression and courtship in *Drosophila* (Goergen et al. 2014). Differential expression of *ube3a* has also been linked to male aggression. Interestingly, the *ube3a* gene is responsible for producing Ubiquitin-protein ligase E3A, an enzyme that aids in the degradation of proteins, which may be adaptive for the protein-rich diet of scale-eaters which exhibit substantial differential expression of metabolism-related genes (McGirr and Martin 2018). Differential expression of *ube3a* has also been linked to variation in levels of aggression in male rats (Kurian et al. 2007; Stoppel 2014). However, snail-eater and generalist pupfish also adhere to a lekking mating system, although there may be quantitative differences in male competition and degree of lekking among species and lake populations (CHM pers. obs.).

The increased female aggression of snail-eaters may also be explained by mating system. Although snail-eaters have been observed mating in the lekking system, not much is known about how their courting behaviors differ from generalists. It is possible that increased levels of female aggression are part of the species’ courting ritual. Alternatively, female aggression may have increased incidentally due to selection for decreased maternal care. Our gene expression data indicates that snail-eaters show increased levels of expression for the *nr2e1* gene compared to generalists. This gene (nuclear receptor subfamily 2, group E, member 1) produces a receptor which has been linked to abnormal brain and eye development, as well as increased aggression and lack of maternal care (Young et al. 2002; Abrahams et al. 2005). It is possible that selection for differing levels of maternal care in snail-eaters, compared to generalists or scale-eaters, also incidentally increased levels of aggression for females. For example, one closely related outgroup to *Cyprinodon, Jordanella floridae*, exhibits paternal care of eggs through pectoral fanning (St Mary et al. 2004).

### Increased aggression due to indirect selection

Alternatively, aggression may have increased via selection on other traits. For example, melanin production and aggression are physiologically linked via the melano-cortin system (Cone 2005; Price et al. 2008). This association has been documented across a wide array of vertebrates and suggests that selection for increased melanin pigmentation in other contexts (e.g. mate choice or camouflage) may incidentally increase aggression (Mcgraw et al. 2003; Ducrest et al. 2008; Price et al. 2008). Indeed, territorial male scale-eating pupfish exhibit jet black breeding coloration, unique among cyprinodontiform fishes, and territorial snail-eating pupfish exhibit the lightest breeding coloration of any *Cyprinodon* species (Martin and Wainwright 2013a). Similarly, selection for morphological traits may also indirectly increase aggression. We found differential gene expression between scale-eater *vs* generalist pupfish in the *gnaq* gene, which is annotated for maternal care (Table 2B). *Gnaq* is one of four Gq class α-subunits and aids in phospholipase C-β – receptor coupling (Offermanns et al. 1998). Silencing this gene produces severe craniofacial defects in mice, especially in the mandible (Offermanns et al. 1998). *C. desquamator* show extreme craniofacial features, including enlarged oral jaws that may be beneficial for scale-eating. Thus, it is intriguing that selection for increased jaw size may have indirectly selected for increased aggression in this species. Given the enlarged oral jaws of most scale-eating species, this may be a general mechanism indirectly contributing to increased aggression in scale-eaters depending on how frequently this genetic pathway is modified.

In conclusion, our data suggest that the aggression hypothesis is not a sufficient explanation for the origin of scale-eating in pupfish. Instead, increased aggression in both specialists indicates that aggression may function in dietary specialization or occupation of a novel niche. Alternatively, increased aggression may be an incidental effect of selection on other ecological or sexual traits. Specifically, the aggression-boldness syndrome, the melanocortin system, selection for increased oral jaw size, or metabolic adaptations for increased intake of protein could all have indirectly increased aggression. Future studies should investigate whether aggression is adaptive for scale- and snail-eating in pupfish.

## Acknowledgements

Funding was provided by the University of North Carolina at Chapel Hill. We thank the BEST commission and Ministry of Agriculture of the Commonwealth of the Bahamas for permission to conduct this research and export specimens. All laboratory behavioral and sampling protocols were approved by the University of North Carolina at Chapel Hill IACUC (protocol# 15-179.0). The Gerace Research Centre and the UNC High Throughput Genomic Sequencing Facility provided logistical support.

## Author contributions

MES and CHM conceptualized the project, MES and JAM collected data and performed analyses, MES wrote the manuscript, and all authors revised the manuscript.

## Data accessibility

All behavioral datasets from this study will be deposited in the Dryad Digital Repository. Transcriptomic raw sequence reads are deposited at the NCBI BioProject database (Title: Craniofacial divergence in Caribbean Pupfishes. Accession: PRJNA391309).

